# Metabolic reprogramming and partial acquisition of cancer stem cell-like phenotype in human umbilical cord-mesenchymal stem cells under hypoxia

**DOI:** 10.64898/2026.03.11.710925

**Authors:** Yoshihiro Kushida, Kana Abe, Yo Oguma

## Abstract

Mesenchymal stem cells (MSCs) cultured in hypoxic conditions have been suggested to have more therapeutic efficacy than those cultured under normoxic conditions, and there is growing interest in using hypoxic MSCs for clinical treatment, particularly human umbilical cord (hUC)-MSCs. We investigated how hUC-MSCs and human bone marrow (hBM)-MSCs change from normoxia to hypoxia (1% O_2_) for 2 weeks of culture. In the growth speed and population doubling time, hUC-MSCs cultured under hypoxia exhibited a significantly higher proliferation rate beyond cancerous cells, such as human glioblastoma and breast cancer cells, while hBM-MSCs did not show a significant difference between normoxia and hypoxia, and were statistically slower than these cancerous cells. Notably, hypoxic hUC-MSCs showed upregulation of genes related to metabolic reprogramming (cholesterol biosynthesis and fatty acid metabolism pathways) and cancer stem cell-like phenotype (factors related to Wnt and Hedgehog signaling pathways, cell proliferation drivers, and apoptosis-resistance), and lesser migration and homing to the traumatic brain injury than normoxic hUC-MSCs after intravenous injection. Thus, whether hUC-MSCs cultured under hypoxia offer clinical benefits and use are safe, given their extremely accelerated proliferation rate and partial cancer stem cell-like traits, requires comprehensive and careful investigation.

## Introduction

Mesenchymal stem cells (MSCs) can be easily obtained as adherent cells from various sources, such as human bone marrow (hBM), adipose tissue, and umbilical cord (hUC), and can be expanded to the clinical scale.^1^ They are non-tumorigenic and have a low risk of tumorigenicity, making them a low hurdle to clinical applications. Particularly, hUC, which can be obtained from medical waste, is one of the most widely used MSC types.^1^ MSCs cultured in hypoxic conditions have been suggested to have greater therapeutic efficacy than those cultured under normoxia, such as growth speed acceleration, maintaining stemness, and enhancing paracrine effect, and there is a great movement to use MSCs cultured under hypoxia for clinical treatment, particularly in hUC-MSCs.^2-6^

Human BM-MSCs and hUC-MSCs both show similar MSC markers and differentiation potential to osteogenic, chondrogenic, and adipogenic lineage cells, whereas hUC-MSCs have higher proliferation capacity and CFU-F (colony-forming capacity) than BM-MSCs.^1^ Human UC-MSCs show distinct immune-related characteristics.^7^ However, it has been reported that MSC properties vary depending on the tissue from which they are derived.^8^ Indeed, tissue oxygen concentrations are different: hBM is 2.5-5 mmHg, and hUC is generally higher, at 25-35 mmHg.^9,10^ Therefore, the characteristics of hBM-MSCs and hUC-MSCs differ in various respects, and their responses to hypoxia would differ among tissue sources. In this study, we investigated how hUC-MSCs and hBM-MSCs change from normoxia to hypoxia, focusing on proliferation rate, changes in gene expression, and migration into damaged tissue in vivo.

## Results and Discussion

Human UC-MSC and hBM-MSCs, both with three batches at passages 6, were cultured under normoxia (21% O_2_) (Normo-hUC-MSCs and Normo-hBM-MSCs) and hypoxia (1% O_2_) (Hypo-hUC-MSCs and Hypo-hBM-MSCs), respectively, for 2 weeks (Fig. 1A). Cell morphology seemed to be largely unchanged between Normo-hBM-MSCs and Hypo-hBM-MSCs, whereas Hypo-hUC-MSCs tended to be smaller and round than Normo-hUC-MSCs (Fig. 1B). U251 human malignant glioma cells and human MCF-7 breast carcinoma cells were set as controls for the proliferation rate assay.

**Figure 1.**
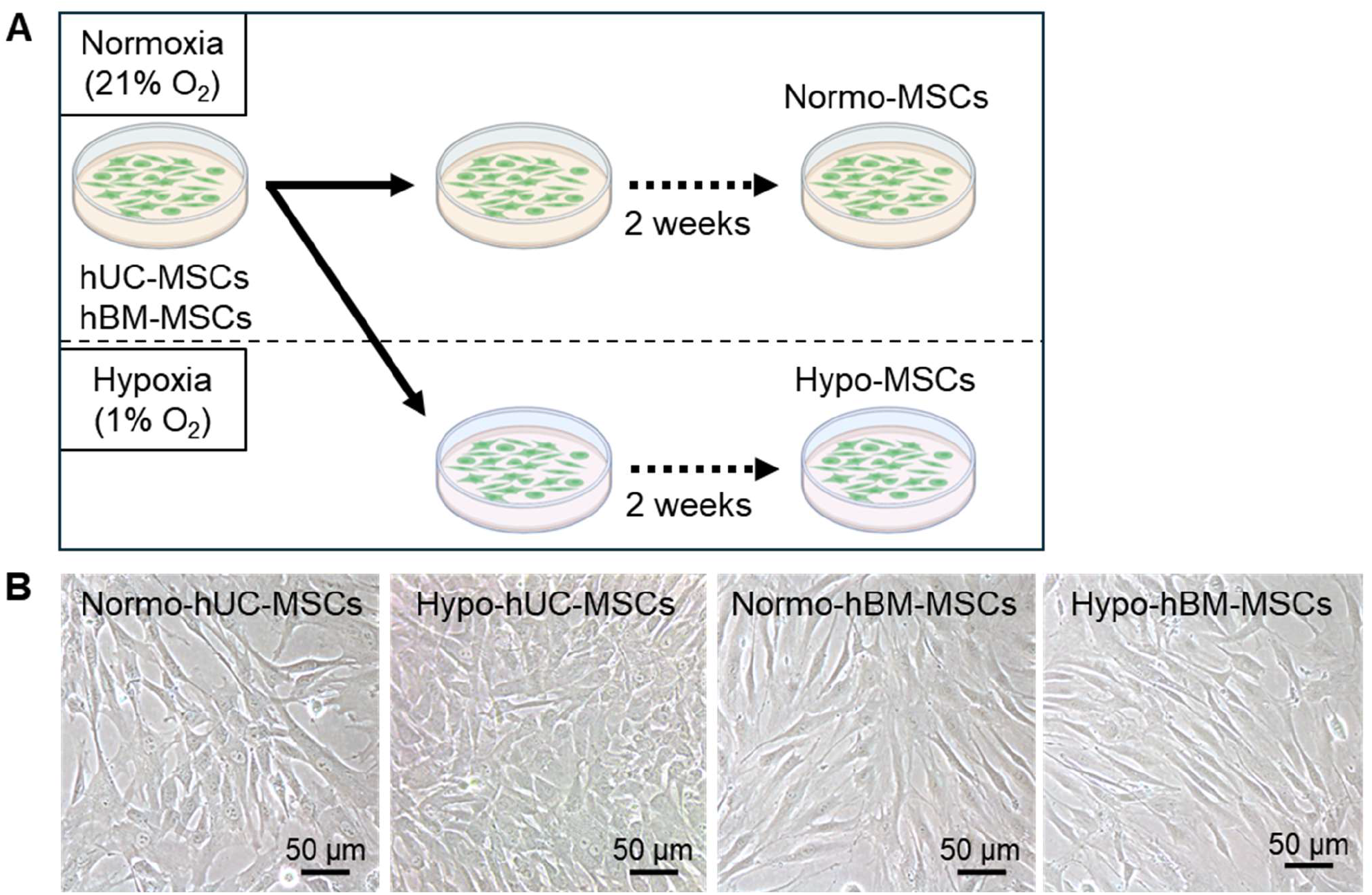
Schematic diagram of the experimental workflow. **(A)** Human UC- (n=3) and hBM-MSCs (n=3) were cultured for 2 weeks under 21% O_2_ normoxic or 1% O_2_ hypoxic conditions. **(B)** Representative phase-contrast images of hUC- and hBM-MSCs under normoxia and hypoxia at 2 weeks.

Despite with the same start seeding density 2,500 cells/cm^2^, Hypo-hUC-MSCs reached 146.4 ± 8.3 x 10^3^ cells by day 7, significantly greater cell number than U251 (57.7 ± 1.5 x 10^3^) (*p* < 0.001) and MCF-7 (81.3 ± 1.5 x 10^3^) (*p* < 0.001), as well as than Normo-hUC-MSCs (36.3 ± 2.3 x 10^3^) (*p* < 0.001), Normo-hBM-MSCs (29.4 ± 0.7 x 10^3^) (*p* < 0.001), and Hypo-hBM-MSCs (27.2 ± 2.6 x 10^3^) (*p* < 0.001) (Fig. 2A). The cell number of Normo-hUC-MSCs, Normo-hBM-MSCs, and Hypo-hBM-MSCs were in the range of 25∼35 x 10^3^ cells, and all of them were smaller cell numbers than those in U251 (*p* < 0.001) and MCF-7 (*p* < 0.001), respectively, with statistical significances (Fig. 2A).

**Figure 2.**
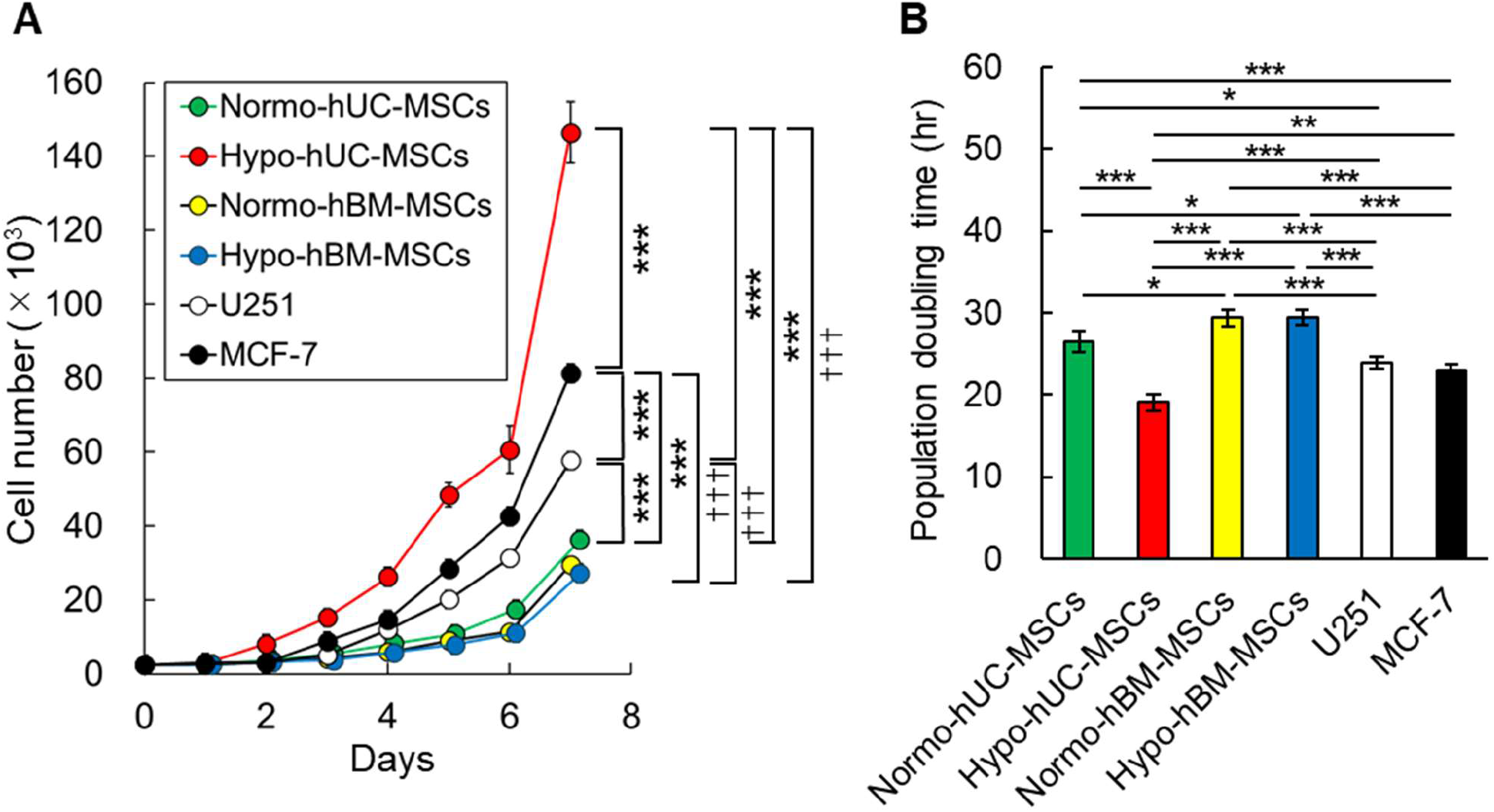
Proliferation of Normo/Hypo-hUC/hBM-MSCs, human glioblastoma, and breast cancer cells. **(A)** Proliferation speed of Normo-hUC-/hBM-MSCs, Hypo-hUC-/hBM-MSCs, malignant glioblastoma cells (U251), and breast carcinoma cells (MCF-7). **(B)** Population doubling time of each group. Data were expressed as the mean ± standard error of the mean (SEM). Symbols (* *p* < 0.05; ** *p* < 0.01; *** *p* < 0.001, ^†††^ *p* < 0.001) indicate significant differences between hUC-MSCs and other cell types (*) and between hBM-MSCs and other cell types (†). One-way analysis of variance (ANOVA) followed by Tukey-Kramer honestly significant difference (HSD) post hoc test was used.

The population doubling time calculated on day 6-7 days in Hypo-hUC-MSCs was 19.1 ± 1.0 hrs, the shortest among all the groups with statistical significances to other five kinds of cells (all *p* < 0.001), and was even shorter than U251 (23.9 ± 0.8 hrs) (*p* < 0.001) and MCF-7 (22.9 ± 0.8 hrs) (*p* < 0.01), suggesting a very strong cell proliferation beyond cancerous cells had been occured (Fig. 2B). In contrast, the population doubling time of Normo-hUC-MSCs (26.0 ± 0.9 hrs) was significantly longer than that of Hypo-hUC-MSCs (*p* < 0.001), and even than that of U251 (*p* < 0.05), and MCF-7 (*p* < 0.001) (Fig. 2B). The longest population doubling time was Normo-hBM-MSCs (31.2 ± 0.8 hrs), followed by Hypo-hBM-MSCs (29.5 ± 0.9 hrs), without statistical significance between them, suggesting that the growth speed of hBM-MSCs was not affected by hypoxia, unlike hUC-MSCs. Both Normo- and Hypo-hBM-MSCs had a longer population doubling time than the other four groups, Normo- (both *p* < 0.05) and Hypo-hUC-MSCs (both *p* < 0.001), U251 (both *p* < 0.001), and MCF-7 (both *p* < 0.001) (Fig. 2B).

The gene expression patterns and levels were compared between Hypo-hUC-MSCs and Hypo-hBM-MSCs to identify differentially expressed genes (DEGs) (Fig. 3A).

**Figure 3.**
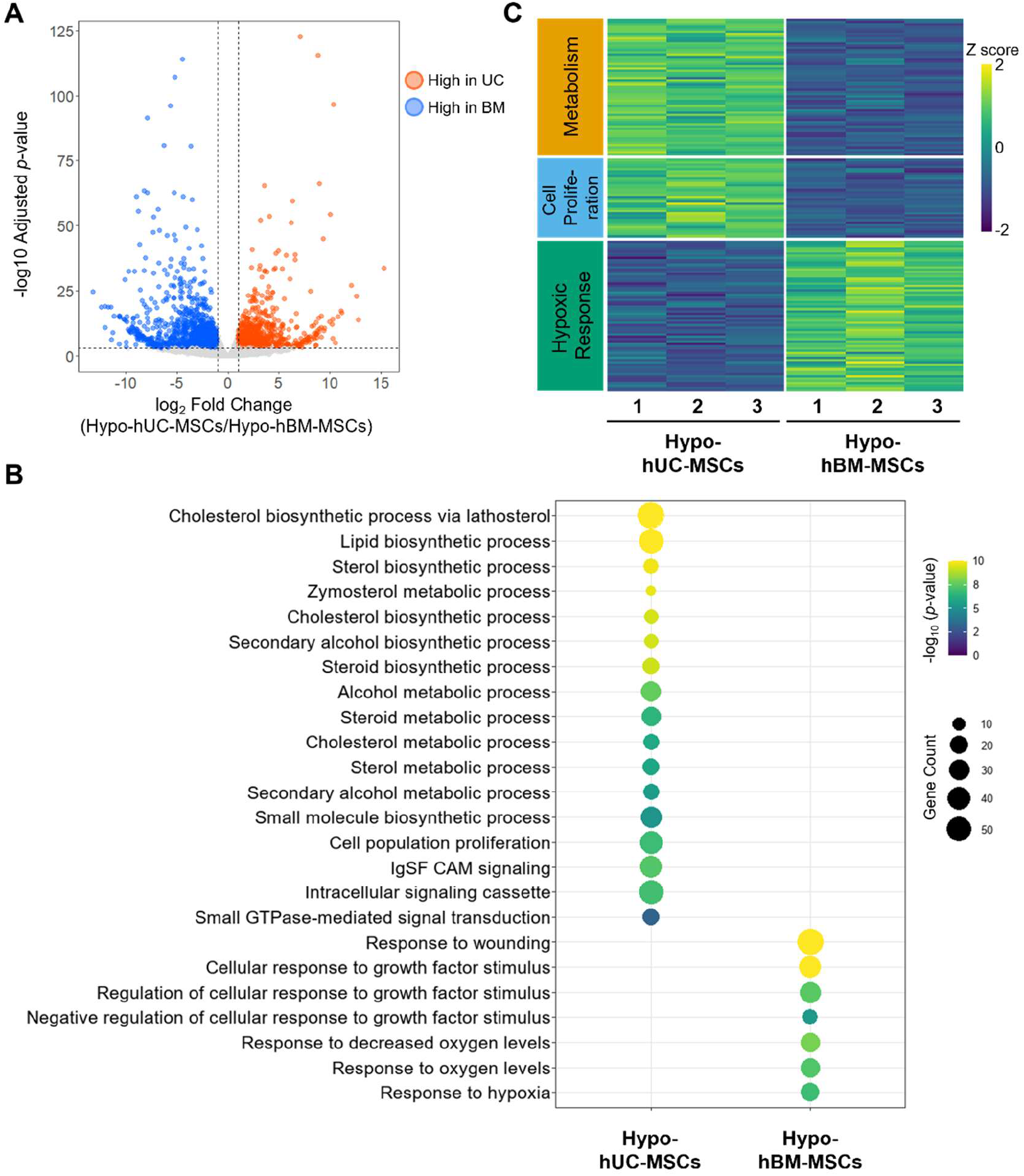
Transcriptomic landscape of cell source-specific responses to hypoxia by RNA sequencing. **(A)** Volcano plots showing differentially expressed genes (DEGs) between Hypo-hUC-MSCs and Hypo-hBM-MSCs. Red and blue dots represent genes significantly up-regulated in Hypo-hUC-MSCs and Hypo-hBM-MSCs, respectively (|log_2_ fold changes| > 1, adjusted *p*-value < 0.001). **(B)** Dot plot of Gene Ontology (GO) enrichment analysis for gene sets significantly up-regulated in Hypo-hUC-MSCs and Hypo-hBM-MSCs. The size of the dots represents the gene count, and the color indicates the significance (-log_10_ [*p*-value]). **(C)** Heatmap of representative genes associated with metabolism, cell proliferation, and hypoxic response. Expression levels are shown as Z-scores of normalized counts. For all RNA-seq data, n=3 independent samples were used per group.

Genes specifically upregulated in Hypo-hUC-MSCs (705 genes) were related to metabolic systems, primarily cholesterol synthesis (cholesterol biosynthetic process, lipid biosynthetic process, sterol biosynthetic process, and zymosterol metabolic process) and cell proliferation (cell population proliferation and Intracellular signaling cassette) (Fig. 3B). On the other hand, genes upregulated in Hypo-hBM-MSCs (500 genes) were associated with canonical hypoxia responses (response to decreased oxygen level and response to hypoxia) and environmental adaptation responses (response to wounding and cellular response to growth factor stimulus) (Fig. 3B). Genes related to metabolism (59 genes), cell proliferation (35 genes) and hypoxic response (77 genes) were extracted and displayed as a heatmap (Fig. 3C).

Notably, when compared Hypo-hUC-MSCs and Hypo-hBM-MSCs in relation to cholesterol biosynthesis pathway, Hypo-hUC-MSCs upregulated isocitrate dehydrogenase 1 (IDH1; converts isocitrate to α-ketoglutarate in an NADP-dependent manner and is also involved in epigenetic regulation) (*p* < 0.01), 3-hydroxy-3-methylglutaryl-CoA synthase 1 (HMGCS1; catalyzes the condensation of acetyl-CoA and acetoacetyl-CoA to form HMG-CoA) (*p* < 0.01), 3-hydroxy-3-methylglutaryl-CoA reductase (HMGCR; a rate-limiting enzyme in the mevalonate pathway) (*p* < 0.01), isopentenyl-diphosphate delta isomerase 1 (IDI1; regulates isomerization of isopentenyl diphosphate) (*p* < 0.01), farnesyl-diphosphate farnesyltransferase 1 (FDFT1; catalyzes the first committed step in sterol biosynthesis) (*p* < 0.001), squalene epoxidase (SQLE; responsible for the late stage of cholesterol synthesis) (*p* < 0.01), transmembrane 7 superfamily member 2 (TM7SF2; participates in the reduction of the sterol C14-double bond) (*p* < 0.01), with statistical significance to Hypo-hBM-MSCs (Fig. 4A). Similarly, in fatty acid metabolism, Hypo-hUC-MSCs upregulated stearoyl-CoA desaturase (SCD; responsible for the desaturation of fatty acids and their recruitment to the lipid synthesis pathway) (*p* < 0.05), ELOVL fatty acid elongase 2 (ELOVL2; catalyzes the elongation of long-chain polyunsaturated fatty acids) (*p* < 0.001), acyl-CoA synthetase long chain family member 3 (ACSL3; mediates the activation of long-chain fatty acids for their incorporation into phospholipids and neutral lipids) (*p* < 0.001), with statistical significance to Hypo-hBM-MSCs (Fig. 4B). These collectively suggested that metabolic reprogramming had been occurred in Hypo-hUC-MSCs.

**Figure 4.**
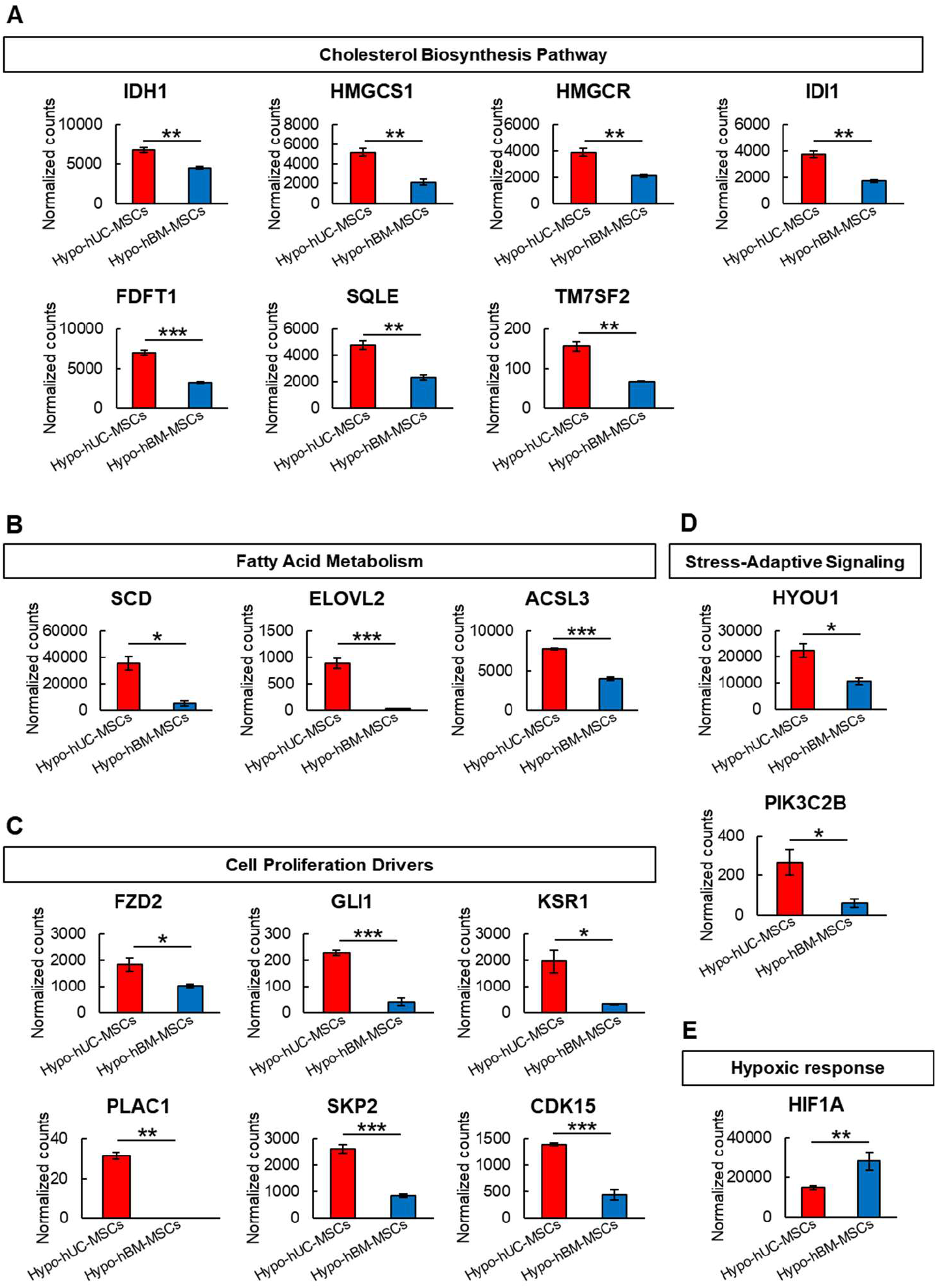
Enhanced metabolic and proliferative signatures in RNA-seq. Representative expression profiles of genes associated with **(A)** Cholesterol biosynthesis pathway (IDH1, HMGCS1, HMGCR, IDI1, FDFT1, SQLE, and TM7SF2), **(B)** Fatty acid metabolism (SCD, ELOVL2, and ACSL3), **(C)** Cell proliferation drivers (FZD2, GLI1, KSR1, PLAC1, SKP2, and CDK15), and **(D)** Stress-adaptive signaling (HYOU1 and PIK3C2B), which are specifically up-regulated in Hypo-hUC-MSCs. **(E)** Representative expression profiles of genes associated with hypoxic response (HIF1A). All expression data are presented as normalized counts to allow for quantitative comparison between groups. Data were expressed as the mean ± SEM (n=3 for each group). * *p* < 0.05; ** *p* < 0.01; *** *p* < 0.001. An unpaired Student’s t-test was used.

Furthermore, expression of cell proliferation drivers, such as frizzled class receptor 2 (FZD2; a Wnt signaling receptor) (*p* < 0.05), glioma-associated oncogene 1 (GLI1; a transcription factor of the Hedgehog signaling pathway) (*p* < 0.001), kinase suppressor of ras1 (KSR1; a RAS signaling regulator) (*p* < 0.05), placenta specific 1 (PLAC1; promotes cell proliferation through activation of the PI3K/Akt pathway) (*p* < 0.01), S-phase kinase-associated protein 2 (SKP2; a G1/S transition promoter) (*p* < 0.001), cyclin dependent kinase 15 (CDK15; regulates cell cycle progression and stress resistance) (*p* < 0.001), as well as stress-adaptation-related genes, such as hypoxia up-regulated 1 (HYOU1; an anti-apoptotic chaperone protein) (*p* < 0.05) and phosphatidylinositol-4-phosphate 3-kinase catalytic subunit type 2 beta (PIK3C2B; generates lipid signaling molecules that facilitate cell proliferation, survival, and membrane trafficking) (*p* < 0.05), were elevated in Hypo-hUC-MSCs than in Hypo-hBM-MSCs with statistical significance (Fig. 4C, 4D).

On the other hand, hypoxia inducible factor 1 subunit alpha (HIF1A; a transcription factor that activates genes for survival and metabolism under hypoxic conditions) was around 2 times higher in Hypo-hBM-MSCs than in Hypo-hUC-MSCs with statistical significance (*p* < 0.01) (Fig. 4E).

Previous reports have shown that BM-MSCs rarely migrate or home to the injury site after intravenous administration.^1^ To examine whether there is a difference between Normo- and Hypo-hUC-MSCs, we intravenously administered 1 × 10^5^ Akaluc-labeled-Normo-hUC-MSCs and -Hypo-hUC-MSCs to a mouse traumatic brain injury (TBI) model 7days after injury. At 3 days post-injection, the in vivo dynamics of the injected cells revealed a faint signal of homing into the TBI in Normo-hUC-MSCs, while the signal was substantially less in Hypo-hUC-MSCs than in Normo-hUC-MSCs (*p* < 0.01) (Fig. 5A, 5B). In contrast, both Normo- and Hypo-hUC-MSCs showed significantly higher accumulation in the lungs, without a statistical difference. No background signals were observed in the brain or lungs of the phosphate buffer saline (PBS)-injected control mice (Fig. 5A).

**Figure 5.**
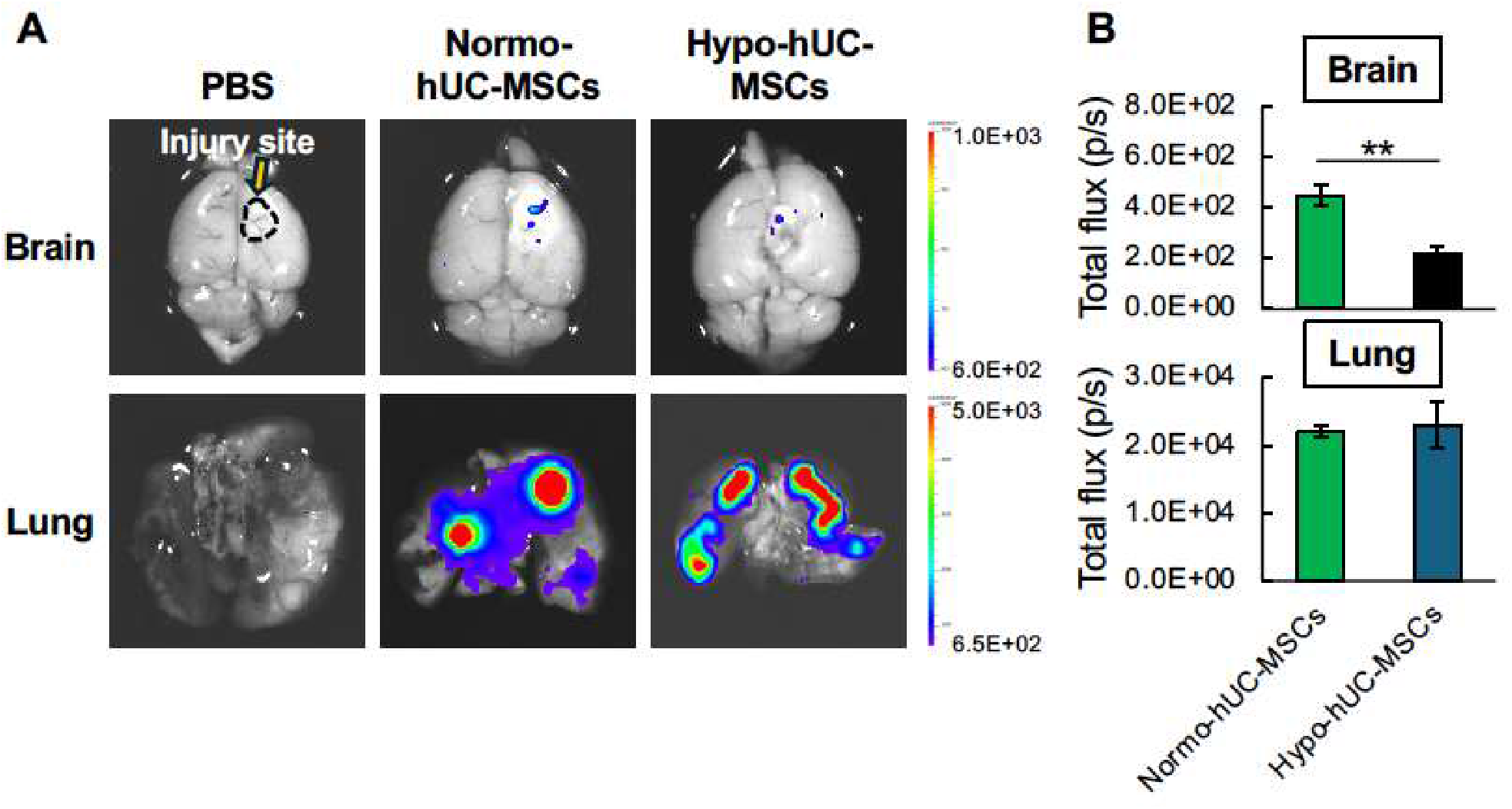
In vivo dynamics of intravenously injected Normo-hUC-MSCs and Hypo-hUC-MSCs in the mouse TBI model. **(A)** Representative ex vivo imaging of the brain and lungs. **(B)** Quantification of photon flux in the brain and lungs. The signal was low in the Normo- and Hypo-hUC-MSCs groups, while Hypo-hUC-MSC group was lower than Normo-hUC-MSC group with statistical significance (*p* < 0.01). Data were expressed as the mean ± SEM (n=3 for each group). ** *p* < 0.01. An unpaired Student’s t-test was used.

MSCs are a heterogeneous population, and their properties vary depending on handling and lot. In this study, three lots were prepared for each category. The following new findings were obtained from this study:

1. Several different responses to hypoxia were observed between hUC-MSCs and hBM-MSCs. First, hUC-MSCs exhibited a significantly higher proliferation rate than cancer cells, such as glioblastoma and breast cancer cells. However, hBM-MSCs showed no significant difference between normoxia and hypoxia, and no significant increase in proliferation was observed.
2. Genes related to metabolic reprogramming (cholesterol biosynthesis and fatty acid metabolism pathways) and cancer stem cell-like phenotype (factors related to Wnt and Hedgehog signaling pathways, cell proliferation drivers, apoptosis-resistance) were upregulated in Hypo-hUC-MSCs.
3. Intravenous injection into the mouse TBI model exhibited that Normo-hUC-MSCs showed limited migration and homing to the damaged TBI tissue, while Hypo-hUC-MSCs showed less migration and homing than Normo-hUC-MSCs, with statistical significance. Therefore, the potential of direct involvement of Hypo-hUC-MSCs at the injury site is unlikely to be enhanced by hypoxic culture.
4. Overall, hUC-MSCs may carry the risk of extremely accelerated proliferation beyond cancerous cells and partial acquisition of cancer stem cell traits. Whether their therapeutic effects are superior to those of Normo-hUC-MSCs needs careful consideration.

The elevation of gene expression associated with an undifferentiated state, such as the Wnt and Hedgehog pathways, as well as factors associated with apoptosis tolerance, such as HYOU1, and increased stress tolerance, as represented by PIK3C2B, are characteristics of a cancer stem cell-like phenotype.^11,12^ Wnt and Hedgehog pathways maintain the self-renewal of cancer stem cells; HYOU1 is associated with metastasis; and PIK3C2B drives cancer progression through metabolic reprogramming.^11-13^

Cancer stem cells are known to reprogram their lipid metabolism to survive and proliferate in the harsh tumor microenvironment.^14,15^ As mentioned above, Hypo-hUC-MSCs also showed a dramatic increase in the expression of genes involved in cholesterol biosynthesis and fatty acid metabolism. This suggests that cell membrane properties are reprogrammed through de novo cholesterol synthesis, such as via HMGCR and SQLE, as well as through the recruitment of fatty acids into the lipogenic pathway via ACSL3 and their desaturation by SCD.^16^ These metabolic reprogramming events go beyond simple component supply and contribute to the reprogramming of highly organized membrane domains, such as lipid rafts. Consequently, under hypoxic conditions, receptors such as FZD2 are over-accumulated in lipid rafts, and the lipid kinase PIK3C2B is activated at the membrane, thereby continuously transmitting potent survival and proliferation signals.^16^ This enhanced downstream signaling might have led to the activation of the key proliferation-regulating transcription factor GLI1, which, in turn, induces the transcription of genes that regulate cell cycle progression, such as SKP2.^17^ Furthermore, PLAC1, which is involved in cell proliferation in normal placental trophoblasts and cancer cells, was specifically elevated in Hypo-hUC-MSCs^18^. This suggests that the hypoxic environment induces the explosive proliferation activity in hUC-MSCs. At the same time, stress-adaptive mechanisms, such as HYOU1 and PIK3C2B, may help prevent hypoxia-induced cell death and support the remarkable proliferation rate of Hypo-hUC-MSCs.^11-13^

On the other hand, genes that were specifically upregulated in Hypo-hBM-MSCs (500 genes), not Hypo-hUC-MSCs, were associated with canonical hypoxia responses (response to decreased oxygen level and response to hypoxia). In this respect, too, the responses of hBM-MSCs and hUC-MSCs to hypoxia differ.

There are risks associated with hUC-MSCs being highly proliferative than tumor cells under hypoxic culture and possibly acquiring, albeit partially, cancer stem cell-like characteristics. However, even under hypoxic conditions, the response of hUC-MSCs is expected to vary with oxygen concentration and culture duration. Whether hUC-MSCs cultured under hypoxia actually offer clinical benefits and whether there are safety concerns due to their extremely accelerated proliferation rate requires comprehensive and careful investigation, including in animal models, in the future.

## Materials and Methods

### Cell culture

Human UC-MSC and hBM-MSCs, both with three batches, were obtained from PromoCell (Heidelberg, Germany) and Lonza (Basel, Switzerland), respectively. The cells were maintained in culture medium comprising α-minimum essential medium (α-MEM) (MilliporeSigma, St Louis, MO, USA) with 10% (vol/vol) fetal bovine serum (FBS) (HyClone, Logan, UT, USA), 1 ng/mL basic fibroblast growth factor (bFGF) (Miltenyi Biotec, Bergisch Gladbach, Germany), 2 mM GlutaMAX (ThermoFisher Scientific, Waltham, MA), and 0.1 mg/mL kanamycin sulfate (Fujifilm, Osaka, Japan) at 37°C in 95% air and 5% CO_2_. Cells from passages 6 were used to prepare cells under normoxic and hypoxic conditions.

LentiX-293 T packaging cells were obtained from Takara Bio (Shiga, Japan). Human breast cancer cell line (MCF-7), and human malignant glioblastoma cell line (U251) were purchased from KAC (Kyoto, Japan). These cells were maintained in 4.5 g/L glucose Dulbecco’s modified Eagle’s medium (DMEM) (ThermoFisher Scientific) supplemented with 10% FBS, 1 mM sodium pyruvate (ThermoFisher Scientific), and 0.1 mg/mL kanamycin sulfate at 37°C in 95% air and 5% CO_2_.

### Culture under normoxic and hypoxic conditions

Normoxic-MSCs and Hypoxic-MSCs, both from hUC and hBM, were prepared by culturing under normoxia (21% O_2_) and hypoxia (1% O_2_), respectively, at 37°C for 2 weeks. For hypoxic culture, the culture plates were placed in a modular incubator chamber (Billup-Rothenburg Inc., Del Mar, CA, USA) and flushed with a gas mixture containing 1% O_2_, 5% CO_2_, and 94% N_2_. The medium was pretreated under hypoxic conditions (1% O_2_) for 1 hour before use. When cells reached 90% confluence, they were subcultured at a 1:3 ratio for hUC-MSCs and a 1:2 ratio for hBM-MSCs.

### Cell proliferation assay

The cells were plated at a density of 2,500 cells/cm^2^ onto a 48-well culture plate. The medium was exchanged every 2 days. Cell counting was conducted manually using a hemocytometer. The cell number was calculated using the average of 3 replicates. The population doubling time (PDT) was calculated using the cell counts from day 6 and 7 during the exponential growth phase, according to the following formula:

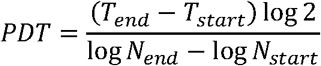

where T represents the incubation time (h) and N represents the cell number.

### RNA sequencing

Total RNAs were extracted using the NucleoSpin RNA Plus XS (Macherey-Nagel, Duren, Germany). The extracted RNA’s purity and concentration were evaluated using an Agilent 2100 bioanalyzer (Agilent Technologies, Palo Alto, CA, USA). The libraries were constructed using NEBNext Poly(A) mRNA Magnetic Isolation Module and NEBNext Ultra RNA Library Prep Kit for Illumina (New England Biolabs, Hitchin, UK). Sequencing was performed on a NovaSeq X Plus (Illumina, San Diego, CA, USA) by Rhelixa (Tokyo, Japan).

Raw reads were subjected to quality control and adapter trimming using fastp (version 0.20.0).^19^ The processed reads were mapped to the human reference genome (GRCh38.94) by HISAT2 (version 2.2.1) with default settings.^20^ Gene counts from the HISAT2 mapping were generated with featureCounts (version 2.0.128) for downstream expression analysis.

DEGs were identified between Hypo-hUC-MSCs and Hypo-hBM-MSCs using DESeq2 (version 1.42.0) in R. The statistical thresholds were set at an absolute log_2_ Fold Change (FC) ≥ 1 and a Benjamini-Hochberg adjusted *p*-value < 0.001.

Functional enrichment analysis was conducted using Metascape (http://metascape.org). For the Hypo-hBM-MSCs group, due to the large number of identified genes, a refined list of the top 500 genes ranked by average expression (baseMean) was curated after applying relaxed criteria (adjusted *p*-value < 0.001, log_2_ FC ≥ -1.25, baseMean > 50) to capture a more comprehensive profile of the environmental response. For the heatmap, Z-scores were calculated from Variance Stabilizing Transformation (VST)-normalized expression values.

### In vivo dynamics of intravenously injected cells in a mouse traumatic brain injury (TBI) model

Eight-week-old male C57BL/6 mice were purchased from Japan SLC Inc. (Shizuoka, Japan). All animals were treated in accordance with the regulations of the Standards for Human Care and Use of Laboratory Animals of Tohoku University. The animal experiments were approved by the Animal Care and Experimentation Committee of Tohoku University Graduate School of Medicine (permission No. 2025MdA-024). Mice were habituated in a pathogen-free environment with a 12-h light–dark cycle for at least 1 week and allowed free access to standard institutional food and tap water. TBI was produced as described previously.^21^ Briefly, anesthesia was induced with 5% isoflurane in N_2_O: O_2_ (70:30) and maintained with 2% isoflurane in N_2_O: O_2_ (70:30). The skull was exposed through a midline skin incision. A freezing brain injury was generated by applying a 3.5-mm-diameter copper 120 cylinder cooled with liquid nitrogen to the right parietal bone 2 mm lateral to the bregma for 180 sec. The scalp incision was sutured, and the animals were allowed to awake from anesthesia. One week after TBI, 1 × 10^5^ Akaluc-labeled-Normo-hUC-MSCs, or -Hypo-hUC-MSCs, each suspended in phosphate-buffered saline (PBS), were injected into the tail vein of mice. Mice receiving an equal volume of PBS served as the control group. Three days post-injection, mice were euthanized by a lethal dose of isoflurane anesthesia. The brain and lungs were removed, immersed in 500 µM AkaLumine-HCl (Fujifilm) in normal saline, and imaged in vivo using an IVIS Spectrum CT2 in vivo imaging system (Revvity, Waltham, MA, USA).^22^ Bioluminescence signals were analyzed as regions of interest using Living Image 4.8.2 software (Revvity). Data are expressed as total photon flux (photons/s).

### Statistical analysis

All data were expressed as mean ± standard error of the mean (SEM). Statistical analysis was conducted using JMP Pro software package (version 18.0.2; JMP Statistical Discovery LLC, Cary, NC). Continuous data were compared by one-way analysis of variance (ANOVA) followed by Tukey-Kramer honestly significant difference (HSD) post hoc test among more than 2 groups. Unpaired Student’s t-test was used to evaluate the significance of differences between 2 groups. A p-value of less than 0.05 was considered significant.

## References

1. Maxson, S., Lopez, E.A., Yoo, D., Danilkovitch-Miagkova, A. & Leroux, M.A. Concise review: role of mesenchymal stem cells in wound repair. Stem Cells Transl Med 1, 142–149 (2012).

2. Huang, Q.M., et al. Long-term hypoxic atmosphere enhances the stemness, immunoregulatory functions, and therapeutic application of human umbilical cord mesenchymal stem cells. Bone Joint Res 13, 764–778 (2024).

3. Lavrentieva, A., Majore, I., Kasper, C. & Hass, R. Effects of hypoxic culture conditions on umbilical cord-derived human mesenchymal stem cells. Cell Commun Signal 8, 18 (2010).

4. Zhu, Z., Zhang, Y., Huang, Z., Hao, H. & Yan, M. Hypoxic culture of umbilical cord mesenchymal stem cell-derived sEVs prompts peripheral nerve injury repair. Front Cell Neurosci 16, 897224 (2022).

5. Zeng, H.L., et al. Hypoxia-mimetic agents inhibit proliferation and alter the morphology of human umbilical cord-derived mesenchymal stem cells. BMC Cell Biol 12, 32 (2011).

6. Jin, H.J., et al. Comparative analysis of human mesenchymal stem cells from bone marrow, adipose tissue, and umbilical cord blood as sources of cell therapy. Int J Mol Sci 14, 17986–18001 (2013).

7. Lee, H.J., et al. Immunologic properties of differentiated and undifferentiated mesenchymal stem cells derived from umbilical cord blood. J Vet Sci 17, 289–297 (2016).

8. Costela-Ruiz, V.J., et al. Different Sources of Mesenchymal Stem Cells for Tissue Regeneration: A Guide to Identifying the Most Favorable One in Orthopedics and Dentistry Applications. Int J Mol Sci 23(2022).

9. Narazaki, A., et al. Determination of the physiological range of oxygen tension in bone marrow monocytes using two-photon phosphorescence lifetime imaging microscopy. Sci Rep 12, 3497 (2022).

10. Hellegers, A.E. & Schruefer, J.J. Nomograms and empirical equations relating oxygen tension, percentage saturation, and pH in maternal and fetal blood. Am J Obstet Gynecol 81, 377–384 (1961).

11. Takebe, N., Harris, P.J., Warren, R.Q. & Ivy, S.P. Targeting cancer stem cells by inhibiting Wnt, Notch, and Hedgehog pathways. Nat Rev Clin Oncol 8, 97–106 (2011).

12. Chou, X., et al. PIK3C2B drives lung cancer progression through coordinating metabolic reprogramming and EMT-mediated metastasis. Biochem Biophys Rep 44, 102380 (2025).

13. Sargiacomo, C. & Klepinin, A. Density Gradient Centrifugation Is an Effective Tool to Isolate Cancer Stem-like Cells from Hypoxic and Normoxia Triple-Negative Breast Cancer Models. Int J Mol Sci 25(2024).

14. Liu, H., Zhang, Z., Song, L., Gao, J. & Liu, Y. Lipid metabolism of cancer stem cells. Oncol Lett 23, 119 (2022).

15. Zhu, Y., et al. CPT1A-mediated MFF succinylation promotes stemness maintenance in ovarian cancer stem cells. Commun Biol 8, 250 (2025).

16. Lee, H., et al. Lipid metabolism in cancer stem cells: reprogramming, mechanisms, crosstalk, and therapeutic approaches. Cell Oncol (Dordr) 48, 1181–1201 (2025).

17. Chan, C.H., Lee, S.W., Wang, J. & Lin, H.K. Regulation of Skp2 expression and activity and its role in cancer progression. ScientificWorldJournal 10, 1001–1015 (2010).

18. Mahmoudian, J., et al. PLAC1: biology and potential application in cancer immunotherapy. Cancer Immunol Immunother 68, 1039–1058 (2019).

19. Chen, S., Zhou, Y., Chen, Y. & Gu, J. fastp: an ultra-fast all-in-one FASTQ preprocessor. Bioinformatics 34, i884–i890 (2018).

20. Kim, D., Paggi, J.M., Park, C., Bennett, C. & Salzberg, S.L. Graph-based genome alignment and genotyping with HISAT2 and HISAT-genotype. Nat Biotechnol 37, 907–915 (2019).

21. Shiraishi, K., et al. Intravenous transplantation of multi-lineage differentiating stress enduring cell promotes functional recovery after traumatic brain injury in mice. Sci Rep (2026).

22. Iwano, S., et al. Single-cell bioluminescence imaging of deep tissue in freely moving animals. Science 359, 935–939 (2018).

